# Genome-wide Population Structure Analyses of Three Minor Millets: Kodo Millet, Little Millet, and Proso Millet

**DOI:** 10.1101/499087

**Authors:** Matthew Johnson, Santosh Deshpande, Mani Vetriventhan, Hari D Upadhyaya, Jason G. Wallace

**Affiliations:** Institute of Plant Breeding, Genetics, and Genomics, University of Georgia, 111 Riverbend Rd. Athens, Georgia; International Crops Research Institute for the Semi-Arid Tropics (ICRISAT), Patancheru - 502324, Telangana, India; Institute of Plant Breeding, Genetics, and Genomics, University of Georgia, 111 Riverbend Rd. Athens, Georgia.

## Abstract

Millets are a diverse group of small-seeded grains that are rich in nutrients but have received relatively little advanced plant breeding research. Millets are important to smallholder farmers in Africa and Asia because of their short growing season, good stress tolerance, and high nutritional content. To advance the study and use of these species, we present a genome-wide marker datasets and population structure analyses for three minor millets: kodo millet (*Paspalum scrobiculatum*), little millet (*Panicum sumatrense*), and proso millet (*Panicum miliaceum*). We generated genome-wide marker data sets for 190 accessions of each species with genotyping-by-sequencing (GBS). After filtering, we retained between 161 and 165 accessions of each species, with 3461, 2245, and 1882 single-nucleotide polymorphisms (SNPs) for kodo, proso, and little millet, respectively. Population genetic analysis revealed 7 putative subpopulations of kodo millet and 8 each of proso millet and little millet. To confirm the accuracy of this genetic data, we used public phenotype data on a subset of these accessions to estimate the heritability of various agronomically relevant phenotypes. Heritability values largely agree with the prior expectation for each phenotype, indicating that these SNPs provide an accurate genome-wide sample of genetic variation. These data represent one of first genome-wide population genetics analyses, and the most extensive, in these species and the first genomic analyses of any sort for little millet and kodo millet. These data will be a valuable resource for researchers and breeders trying to improve these crops for smallholder farmers.

## Introduction

It is estimated that there are up to 7,000 cultivated crop species in the world (Khoshbakht and Hammer 2008), yet major breeding and research efforts have focused on just a small number of these (Hammer et al., 2001). Many crops that have been ignored by modern research are still essential to the local communities that have relied on them for thousands of years, and they have potential for diversifying cropping systems around the world (Naylor et al., 2004). These crops are also genetic resources to increase global food security as climate changes and resources (land, water, fertilizers) become more limited. The ability to generate inexpensive, genome-wide data can quickly bring some of these crops into the modern genomics era (Varshney et al 2012).

Millets are a diverse group of small-seeded grains that have been largely overlooked by modern genetics research. Although pearl millet (*Pennisetum glaucum* syn. *Cenchrus americanus*) and foxtail millet (*Setaria italica*) both have complete genome sequences and well-established germplasm resources (Varshney et al. 2017; Zhang et al 2012; Prasad 2017; Sehgal et al 2015), most other millets have few if any resources available (Goron & Raizada 2015). These crops are often important for smallholder farmers, especially in southeast Asia and Sub-Saharan Africa (FAO; http://www.fao.org/docrep/W1808E/w1808e0e.htm).

Proso millet (*Panicum miliaceum* L.), little millet (*Panicum sumatrense*), and kodo millet (*Paspalum scrobiculatum* L.) are three minor millets with very few modern genetic resources (Table 1). Various germplasm repositories maintain collections of these species (Goron & Raizada 2015); the current study focuses on those held by the International Crops Research Institute for the Semi-Arid Tropics (ICRISAT), which maintains collections with 849, 473, and 665 accessions of proso, little, and kodo millet, respectively. These collections have been assessed for morphological and agronomic traits, and representative core collections have been created for each of them (Upadhyaya et al., 2014, 2011). These millets are hardy C_4_ grasses (Upadhyaya et al., 2014, and Brown, 1999) with nutritional content on par with or superior to the major grains (Vetriventhan and Upadhyaya, 2018; Mengesha 1966, Saleh et al., 2013, Kalinova and Moudry 2006).

### Proso Millet

Proso millet is believed to have been independently domesticated ~10,000 years ago in three locations: Northwest China (Bettinger et al., 2007, 2010a,b), Central China (Lu et al., 2009), and Inner Mongolia (Zhao, 2005). It is the third-oldest cultivated cereal after wheat and barley (Habiyaremye et al., 2016; Upadhyaya et al., 2011). Proso millet is valued for its low water requirements (330 mm) and short growing season (60 days) (Habiyaremye et al., 2016; Shanahan et al., 1988). Proso millet varieties are classified into five races based on inflorescence morphology: miliaceum, patentissimum, contractum, compactum, and ovatum (de Wet JMJ, 1986). Proso millet is tetraploid (2n = 4x = 36) (Saha et al., 2016), with evidence suggesting it is an allotetraploid (Habiyaremye et al., 2016). Proso millet has historically be the most widely grown of the three millets studied here, with cultivation concentrated in the former Soviet Union and India (Roshevits, 1980).

Proso millet genetic diversity has been investigated with a variety of genetic markers, most of them at a very small scale (<100 markers; reviewed in Habiyaremye et al 2017). Recently, however, the most extensive genetic analysis in proso millet identified over 400,000 SNP markers and 35,000 SSRs from the transcriptomes of two proso millet accessions (Yue et al 2016). Population-level analyses were made by Rajput et al. (2016) using 100 SSRs and 90 proso millet accessions, it was found that there were some connections between genetic clustering and geographic origin.

### Kodo Millet

Kodo millet was domesticated in India around 3,000 years ago, and India has historically been the major center of cultivation (de Wet et al., 1982). Kodo millet accessions have been classified into three races based on panicle morphology: regularis, irregularis, and variabilis (de Wet et al., 1983; Prasad Rao et al., 1993). Kodo millet is tetraploid (2n = 4x = 40) (Saha et al., 2016), and it is valued for its ability to produce consistently in hot, drought-prone arid and semiarid land (Dwivedi et al., 2012). A few sets of molecular markers have been developed for kodo millet based on RAPD markers (M’Ribu & Hilu 1996), gene-specific primer sets (Kushwaha et al 2015), and semi-targeted PCR amplification (Yadav et al. 2016); the latter study concluded that the 96 accessions they used could be divided into four groups that showed little connection between geographic region and genetic relationship of the accessions. There have been no truly genome-wide datasets on this species before now.

### Little Millet

Little millet was domesticated 5,000 years ago in India (de Wet et al., 1982). It has historically been grown mainly in India, Mayanmar, Nepal, and Sri Lanka (Prasad Rao et al., 1993). Little millet accessions have been classified into 2 races based on panicle morphology: nana and robusta, with two subraces per race (laxa and erecta for nana, and laxa and compacta for robusta) (de Wet et al., 1983a; Prasad Rao et al., 1993). Little millet is tetraploid (2n = 4x = 36) (Saha et al., 2016). Like kodo millet, little millet can give consistent yields on marginal lands in drought-prone arid and semi-arid regions, and it is an important crop for regional food stability (Dwivedi et al., 2012). Little millet is arguably the least studied of these three millets, and we are unaware of any molecular markers developed for it outside of specific single genes (Goron & Raizada 2015) and a small set of RAPD markers whose details were not described (M. S. Swaminathan Research Foundation 2000).

### Expanding genetic resources

As mentioned above, the genetic and genomic resources of proso millet, kodo millet, and little millet are very limited (Saha et al., 2016; Goron & Raizada 2015). The main focus of the resources have been centered on the accessions stored in gene banks, expression sequence tags (ESTs), and complete coding sequences (CDSs).

Core collections for each of these species were created several years ago, consisting of ~10% of ICRISAT’s collection for each species (Upadhyaya et al., 2011,2014). These collections have been assessed for morpho-agronomic traits but no genetic data has been available for them. Our goal in this project was to generate genome-wide marker data on each of these species to enable population genetic analysis and empower breeders and researchers to make more informed decisions about germplasm selection and representation. We genotyped 190 accessions of each species, including the entire core collections of each, using genotyping-by-sequencing (Elshire et al. 2011) to generate the most comprehensive population genetics resource for each of these species to date. These data dramatically expand the genomic resources available to each of these crops and will help make better use of them moving forward.

**Table 1.**
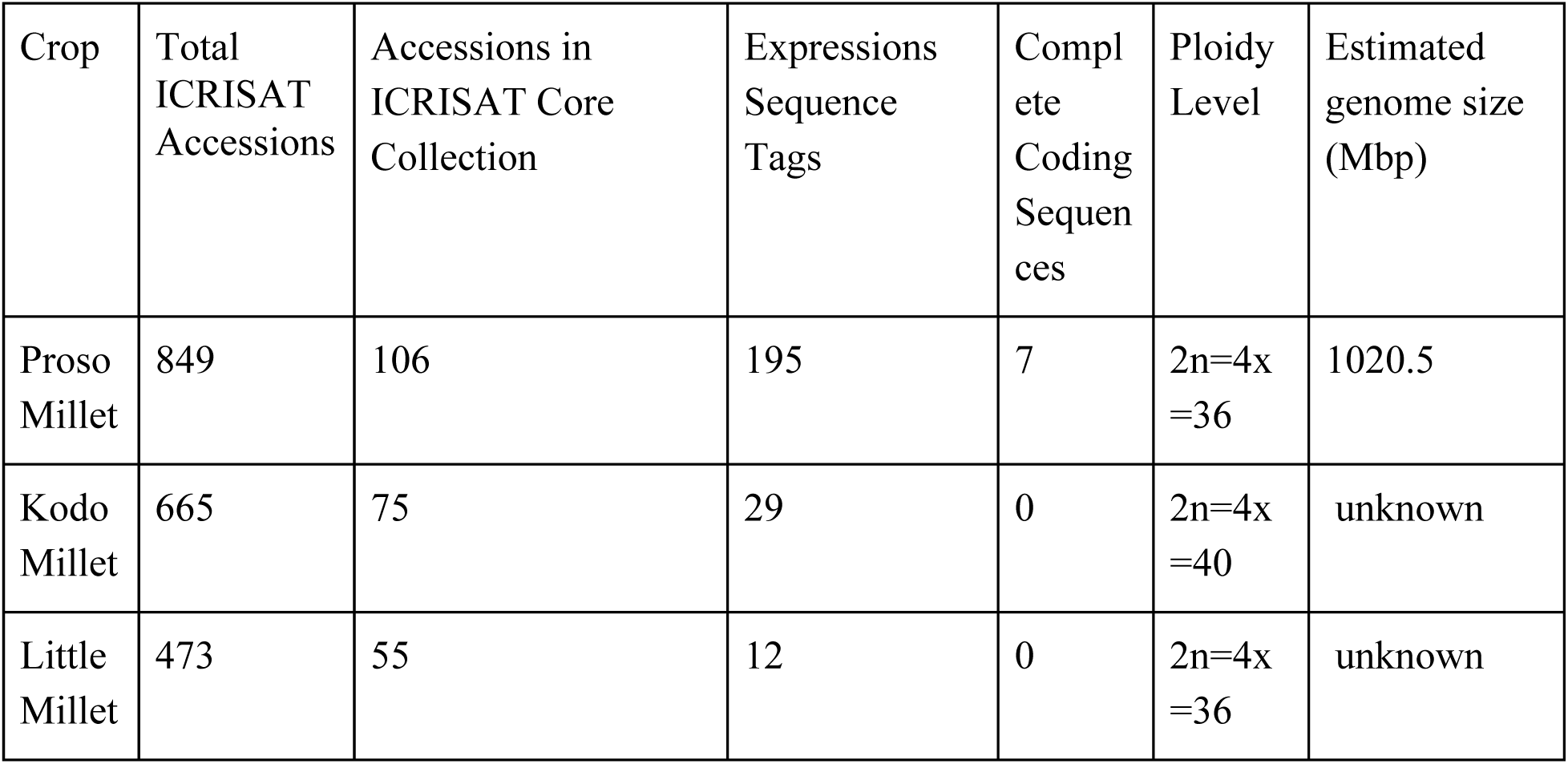
Genetic resources including: Total ICRISAT Accessions (http://genebank.icrisat.org/Default, accessed November 2018), Proso millet core collection (Upadhyaya et al., 2011, Kodo and little millet (Upadhyaya et al., 2014), ploidy information (Upadhyaya et al., 2011 & 2014), and estimated genome size for proso millet (Kubesova et al., 2010)

| Crop | Total ICRISAT Accessions | Accessions in ICRISAT Core Collection | Expressions Sequence Tags | Complete Coding Sequences | Ploidy Level | Estimated genome size (Mbp) |
| --- | --- | --- | --- | --- | --- | --- |
| Proso Millet | 849 | 106 | 195 | 7 | 2n=4x=36 | 1020.5 |
| Kodo Millet | 665 | 75 | 29 | 0 | 2n=4x=40 | unknown |
| Little Millet | 473 | 55 | 12 | 0 | 2n=4x=36 | unknown |

## Materials and Methods

### Plant materials

One-hundred ninety accessions (supplemental folder 3_PC_race_anlysis) were taken from the ICRISAT genebank for each millet species. These included the accessions from the core collection for proso millet, kodo millet and little millet (Upadhyaya et al., 2011, 2014). The remaining samples were chosen to broadly sample the available diversity based on the cluster information used to create the core collections.

### DNA Extraction and sequencing

Seedlings of each accession were grown at the ICRISAT research station in Patancheru, India, in 2015. DNA was extracted by the modified CTAB method (Mace et al., 2003), lyophilized, and shipped to the Genomic Diversity Facility at Cornell University for genotyping-by-sequencing (GBS) (Elshire et al., 2011). GBS library preparation followed standard methods (Wallace & Mitchell 2017) using the PstI restriction enzyme. For kodo millet, 190 samples plus 2 blanks were multiplexed into a single lane for sequencing on an Illumina HiSeq 2500 with single-end 100 bp sequencing. For little and proso millet the procedure was similar, except that samples were multiplexed into 2 lanes of 95 samples plus 1 blank. The raw sequence data for these samples is available at the Sequence Read Archive, accessions PRJNA494158.

### SNP calling

Single-nucleotide polymorphisms (SNPs) were called from raw sequencing data using the TASSEL (Bradbury et al., 2007) GBS v2 SNP pipeline. Since this pipeline uses alignment to a reference genome and none of these species has such a reference, we included slight alterations to make use of the UNEAK filter (Lu et al., 2013) for reference-free alignment of tags to each other. All code for this analysis is available in supplemental folder 7_Code and at https://github.com/wallacelab/2018_minor_millets

### Phylogeny

Phylogenetic networks were constructed using the NeighborNet method (Bryant & Moulton, 2004) in SplitsTree4 V4.14.4 (Huson & Bryant, 2006). Genotype data was converted to SplitsTree-compatible NEXUS files by first using TASSEL version 5.2.29 (Bradbury et al., 2007) to export as a PHYLIP (interleaved) format, which was then converted to NEXUS format using Alter (Glez-Peña et al., 2010) (http://www.sing-group.org/ALTER/). The resulting files were manually edited to change the data type to “dna” and replace all colons (:) with underscores (_), at which point the files were loaded into SplitsTree for network creation.

### Population Structure determination

Population structure analysis was performed with FastStructure v1.0 (Anil et al., 2014), with the number of potential populations (k) varying from 1 to 15. The optimum population size was determined by chooseK and results were visualized with Distruct, both parts of the FastStructure software package. Default parameters were used for all programs. Samples were assigned to a population if they had at least 60% membership in that population.

### Plotting Accessions by Geographic Data

Geographic data on kodo millet and little millet was plotted in python using basemap (Hunter 2007) and the known GPS coordinates, then colored by subpopulation. Since proso millet had only the country of origin for some accessions, its map was created manually in Inkscape (https://inkscape.org/release/0.91/, 2015) by placing dots (colored by subpopulation) in the country of origin for each accession.

### Principal coordinate analysis

Genetic principal coordinates were calculated using multidimensional scaling (MDS) analysis in TASSEL V5.2.29 (Bradbury et al., 2007).

### Heritability analysis

Phenotype data for heritability analysis was taken from public data on the kodo and little millet core collections (Upadhyaya et al., 2014). Phenotypes were fit as part of a mixed linear model in TASSEL (Bradbury et al., 2007) with a kinship matrix as the only covariate, using default parameters. Narrow-sense heritability (h^2^) was estimated as the ratio of genetic variance to total variance in the model.

### Software

The following software and packages were used as part of this analysis:

•: SplitsTree4: V4.14.4: Huson & Bryant, 2006
•: Faststructure: 1.0: https://rajanil.github.io/fastStructure/ Raj et al., 2014
•: TASSEL 5.2.29: http://www.maizegenetics.net/tassel Bradbury et al., 2007
•: PLINK 2.0: http://zzz.bwh.harvard.edu/plink/ Purcell 2007
•: Inkscape 0.91: https://inkscape.org/release/0.91/ 2015
•: GNU Parallel: Tange 2011
•: Matplotlib 2.2.2: Hunter 2007
•: Pandas: McKinney 2010
•: NumPy: Oliphant 2006
•: Python 3.5.2

## Results and Discussion

### Genome-wide marker sets

Genotype data was generated using GBS (Elshire et al. 2011) using PstI restriction digestion. This resulted in ~12–14,000 raw SNPs per species (Table 2), although many of these are probably artifacts due to misalignments. Low-quality SNPs were removed by filtering the raw genotype data to remove sites with >50% missing data, minor allele frequency <5%, and >20–25% heterozygosity. (The heterozygosity filter removes false SNPs due to paralogs misaligning in polyploid species (Wallace et al 2014)). Samples with >50% missing data across all remaining sites were also filtered out. The final, filtered genotype data (supplemental folder 6_HapMaps) consists of 3,461 SNPs across 165 accessions of kodo millet, 2,245 SNPs across 165 accessions of little millet, and 1,882 SNPs across 161 accessions of proso millet. (Table 2)

**Table 2:** Number of SNPs and accessions before and after filtering

| **Species** | **Raw SNPs** | **Filtered SNPs** | **Raw accessions** | **Filtered accessions** |
| --- | --- | --- | --- | --- |
| Kodo Millet | 12995 | 3461 | 190 | 165 |
| Little Millet | 14473 | 2245 | 190 | 165 |
| Proso Millet | 12839 | 1882 | 190 | 161 |

### Proso millet

Population structure analysis of proso millet with fastStructure (Raj et al., 2014) grouped the 161 post-filtering samples into 8 putative subpopulations (Figure 2). Some of the subpopulations consist almost entirely of “pure” individuals of that subpopulation, implying strong separation from the other subpopulations (e.g., groups 7 and 8), while others (e.g., groups 2 through 5) show significant admixture among the subpopulations. We observed that small changes in fastStructure parameters would significantly alter the division and number of subpopulations in this admixed group (data not shown), further indicating that the divisions among the highly admixed subpopulations are weak and should be interpreted with caution.

A phylogenetic network of these samples (Figure 1) mirrors this, where most of the subpopulations group together, although subpopulations 6 and 8 are strongly separated from the rest. Divisions among the main group of populations appear weak, as evidenced by the large number of alternative splits (“webbing”) in the phylogenetic web and the fact that two populations (2 and 4) are split into 2–3 groups across the phylogeny. The genetic principal coordinates of these samples (Figure 2) indicate a similar pattern, where most samples cluster together but with subpopulations 6 and 8 distinct.

While the race data was not available for proso millet in this study there is strong evidence that race is not a good indicator of genetic relatedness among accessions (Vetriventhan & Upadhyaya 2018). Existing core collections were designed with the races of the species being a central focus. Core collections could likely be improved by using population structure and genomic data to improve the genetic diversity of the collections.

The plotting of the 109 accessions with known geographic location supports the population analysis (see supplemental folder 2_maps).

### Kodo millet

Population structure analysis places the 165 kodo millet samples into 7 putative subpopulations (Figures 2). Most accessions cluster together, but 2 subpopulations (5 and 7) are strongly separated from the rest Figures (1,2). The separation of subpopulation 7 from the other samples drives the largest genetic differences in this species, with 67.5% of the genetic variation along this axis (PC1 in Figure 2).

These patterns of population structure do not correlate with existing race designations (supplemental folder 3_PC_race_analysis), implying that existing race designations do not strongly correlate to genetic groupings. Similar to proso millet, this implies that core collections of kodo millet could be improved by using population and genomic data instead of race designation.

Only 17 of the kodo millet accessions had known geographic origins, all of them in India. Plotting these origins geographically supports the population analysis by showing clustering of population 4 (see supplemental folder 2_maps), although more samples would be needed to completely confirm this interpretation.

### Little millet

The 165 filtered accessions of little millet were grouped into 8 putative subpopulations by fastStructure (Figure 2). Little millet appears to have weaker population structure than the other two species (compare the variance explained by each principal coordinate, Figure 2). Subpopulation 5 is relatively well separated from the others, followed by subpopulations 6 and 7, and then the other subpopulations being relatively close together. (These subpopulation designations would shift around with minor changes in fastStructure parameters, implying they are very weakly separated; data not shown.) Similar to the other two species, little millet population structure did not correlated with existing race designations (supplemental folder 3_PC_race_analysis). Plotting the 29 accessions with known geographic location supports the population analysis by showing distinct clusterings of populations 4 and 5 (see supplemental folder 2_maps).

**Figure 1.**
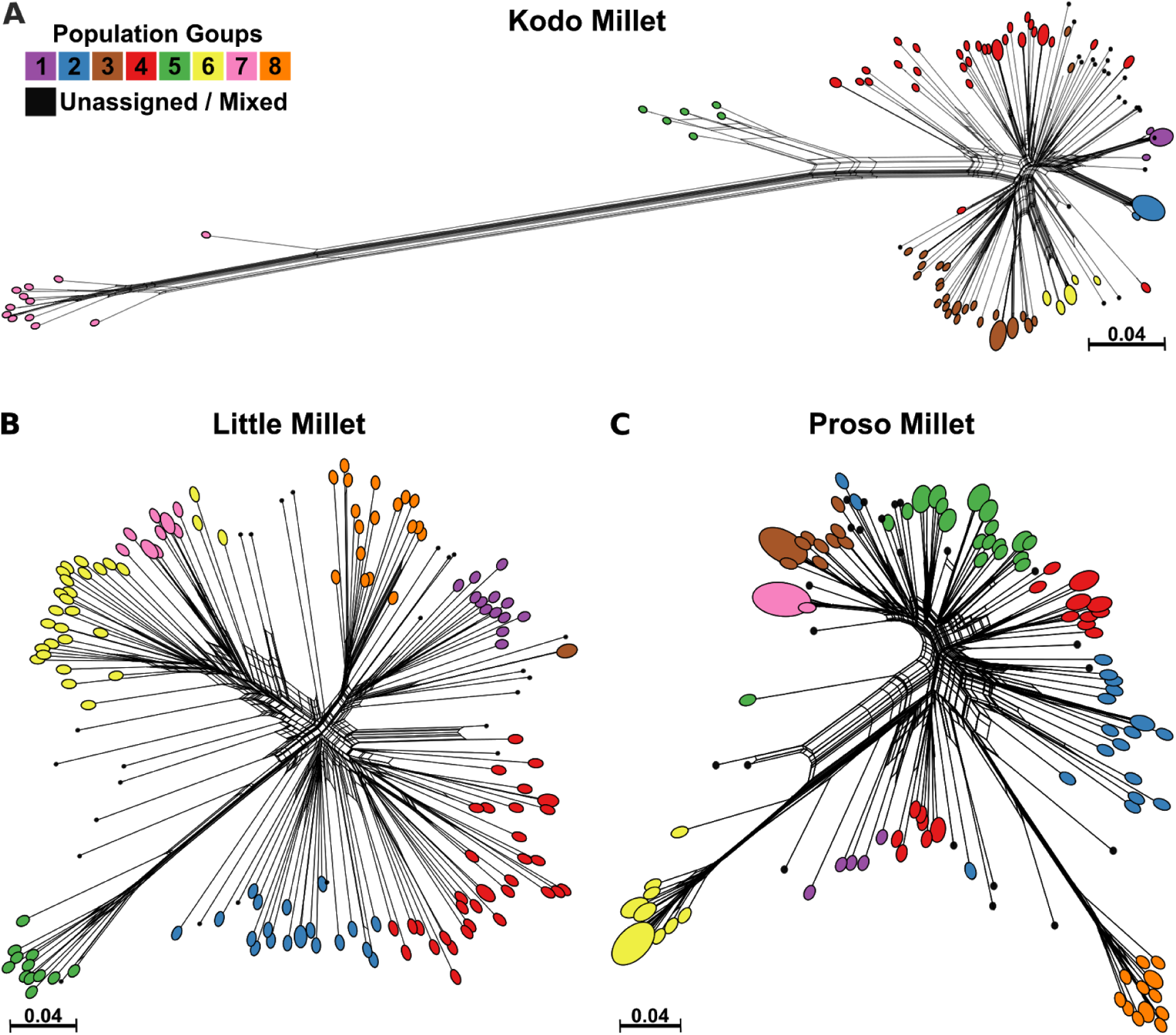
Phylogenetic trees for Kodo Millet (A), Little Millet (B), and Proso Millet (C), colored by population calls from fastStructure for accessions that fit into a population with at least 60% fit. Each accession has one line that ends in a point and has branching to show alternative branching options. Where multiple points of the same population grouped closely together a single oval with a size proportional to the number of accession it encompases was used.

**Figure 2.**
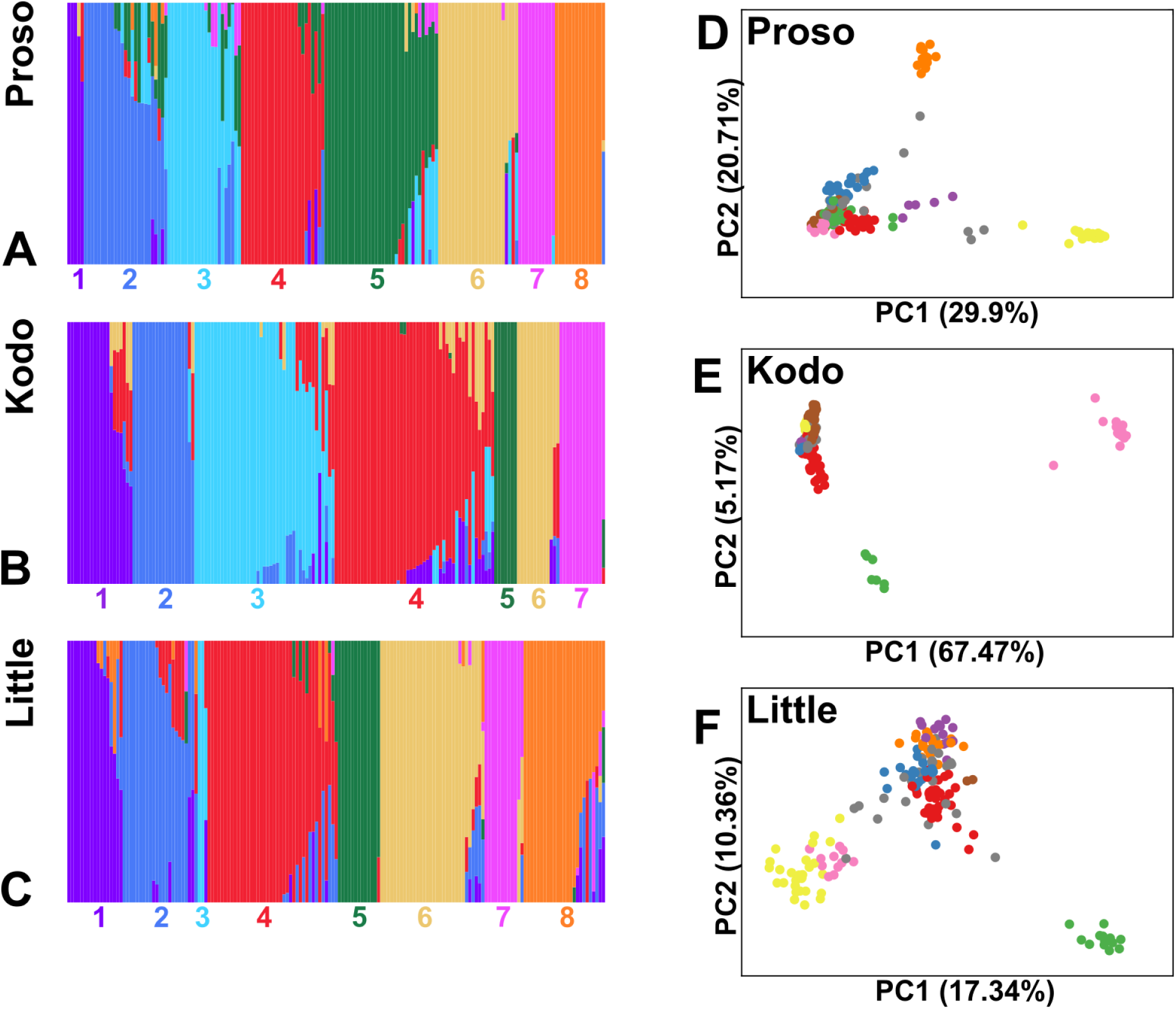
A-C Visual representation of how well individual accessions fit into putative populations from distruct, a part of the fastStructure program. The vertical bars each represent a single accession, and the colors correspond to populations as listed on the bottom of each subfigure (Proso-A, Kodo-B, and Little-C). D-F Principle coordinate analysis based off of genetic distances and colored by putative populations for Proso Millet (A), Kodo Millet (B), and Little Millet (C).

### Validation of Genotype Data via Estimated Heritability

To validate these genetic data, we used public phenotype data on the kodo millet and little millet core collections (Upadhyaya et al., 2014), and proso millet phenotypic data that was provided by ICRISAT. These data were used to estimate narrow-sense heritability (h^2^) for flowering time and plant height (Tables 3). Our expectation was that true genotype data should result in moderate to high heritability values for some phenotypes, especially ones (such as flowering time and plant height) that are known to have a strong genetic component.

**Table 3:** Estimated heritabilities for little millet

| Species | Trait | Heritability |
| --- | --- | --- |
| Little | Days to 50% flowering | 0.807 |
| Little | Plant height (cm) | 0.664 |
| Proso | Days to 50% flowering | 0.69 |
| Proso | Plant height (cm) | 0.615 |
| Kodo | Days to 50% flowering | 0.505 |
| Kodo | Plant height (cm) | 0.293 |

Heritability was estimated using a mixed linear model in TASSEL (see Methods) to obtain estimated variance components. Little millet traits exhibited heritability from 0.209 to 0.807, while kodo millet traits were slightly lower, ranging from 0.0 to 0.505 (supplemental folder 8_Heritability). In all three species flowering time and plant height were some of the most heritable traits (Table 3). These traits are all known to be under strong genetic control in other grass crops (e.g., Buckler et al 2009; Peiffer et al 2014; Mace et al 2013; Morris et al 2013; Ma et al 2016; Alqudah et al 2013). Since poor-quality SNPs should show little to no relationship with phenotype, these results imply that the SNP datasets we have generated accurately represent the genetic variation within these populations and can be used for real-world breeding applications.

## Conclusions

The results from this study represent the first genome-wide analyses for proso millet, kodo millet, and little millets, three “orphan” crops that are important for food security in developing nations. We have identified thousands of SNPs in each of these species that accurately capture the population structure of each, as indicated by the geographic correlations and estimates of narrow-sense heritability. Our analyses can be used as a foundation for further exploration into the genetics of these species, including selecting appropriate breeding materials and identifying priority populations for further collection and curation.

Existing core collections were designed with the races of the species being a central focus. The evidence strongly implies that race is not a good determiner of genetic relatedness, and as such the core collections could likely be improved by using modern genomic data to improve the genetic diversity of the collections.

For both major and minor crops, obtaining genetic data is now (almost) trivial, even in species with highly complex genomes and no prior history of genetic analysis. As the price of DNA sequencing continues to drop, more and more orphan species will have genotype data available. The major question going forward will be how to best deploy these data to benefit breeders and growers. Given that the price of phenotyping is often the limiting factor in many studies (Cobb et al., 2013), finding ways to deploy genomic prediction and/or high-throughput phenotyping for orphan crops will likely be the next major step to democratize modern genomics for the developing world.

## Supporting information

## Author Contributions

Santosh Deshpande, Mani Vetriventhan, Hari Upadhyaya grew the plants and extracted the DNA. Jason Wallace called the SNPs, and Matthew Johnson cleaned and analyzed the data. Matthew Johnson wrote the first draft of the paper and all authors contributed to edits.

## Abbreviations

CDS: coding sequences
ESTs: expressed sequence tags
FAO: Food and Agriculture Organization
GBS: Genotyping-by-Sequencing
ICRISAT: International Crops Research Institute for the Semi-Arid Tropics
MDS: multidimensional scaling
PCR: Polymerase chain reaction
RAPD: Random Amplification of Polymorphic DNA
SNP: single nucleotide polymorphism

