## Supplementary material for "Genome-wide Population Structure Analyses of Three Minor Millets: Kodo Millet, Little Millet, and Proso Millet": 0_Minor_Millet_Filter__and_population_calling_readme

To get the filter working was awful because i didnt properly document everything the first time, here is the step by step

Take the Raw hapmaps and get the GENOTYPE SUMMARY 
Using that raw genotype summary use the following script to get lists of:
Minor Allele Frequency site, Proportion Missing Sites, and Proportion Heterozygous sites

	#Set your working directory to where every you downloaded your files too
	workdir='/home/mjohnson/Desktop/Minor_millet_code_for_publication/'
	#this code takes the site summaries, which are made in tassel from the hapmaps, and extracts site names that fall in
	#parameters found below
	def Filter_function(folder,file_name,collum_name,min_number,max_number,new_file_name):
	    f=open(workdir + folder + "/"  + file_name , "r")
	    column=-1
	    values=list()
	    for line in f:
		x=line.strip("\n").split("\t")
		if column==-1:
		    column=x.index(collum_name)
		    column2=x.index("Site Name")
		y=(x[column])
		q=(x[column2])
		values.append([y,q])
	    filtered=list()
	    for score,name in values:
		if score!=collum_name and float(score) >= min_number and float(score)<= max_number:
		    filtered.append(name+'\n')
		    #print("Site",name,"passed with score",score)
	#         else:
	#             print("Site",name,"failed with score",score)
	    
	    o=filtered
	    h=open (workdir + new_file_name+ ".txt", "w")        
	    h.writelines(o)
	    h.close()
	Filter_function("Site_summaries","Kodo_raw_geno_sum.txt","Proportion Missing",0.0,0.5,"kodo_miss_lt_0.5")
	Filter_function("Site_summaries","Kodo_raw_geno_sum.txt","Proportion Heterozygous",0.0,0.25,"kodo_hetero_lt_0.25")
	Filter_function("Site_summaries","Kodo_raw_geno_sum.txt","Minor Allele Frequency",0.05,0.5,"kodo_min_al_freq_gt_0.05")
	Filter_function("Site_summaries","Little_raw_geno_sum.txt","Proportion Missing",0.0,0.5,"little_miss_lt_0.5")
	Filter_function("Site_summaries","Little_raw_geno_sum.txt","Proportion Heterozygous",0.0,0.25,"little_hetero_lt_0.25")
	Filter_function("Site_summaries","Little_raw_geno_sum.txt","Minor Allele Frequency",0.05,0.5,"little_min_al_freq_gt_0.05")
	Filter_function("Site_summaries","Proso_raw_geno_sum.txt","Proportion Missing",0.0,0.5,"proso_miss_lt_0.5")
	Filter_function("Site_summaries","Proso_raw_geno_sum.txt","Proportion Heterozygous",0.0,0.2,"proso_hetero_lt_0.2")
	Filter_function("Site_summaries","Proso_raw_geno_sum.txt","Minor Allele Frequency",0.05,0.5,"proso_min_al_freq_gt_0.05")


Next Use the following script to get rid of duplicate sites in the three previous site files:
	#Set your working directory to where every you downloaded your files too
	workdir='/home/mjohnson/Desktop/Minor_millet_code_for_publication/'
	#This uses pandas to delete duplicate rows
	def Dup_del(file, new_name):
	    import pandas as pd
	    hmp_df=pd.read_table(workdir + file + ".txt")
	    y=hmp_df.drop_duplicates(keep=False)

	    y.to_csv(workdir + new_name + '.txt', index=False, sep="\t")
	    print('Done')
	    
	Dup_del('kodo_miss_lt_0.5','No_dup_kodo_miss_lt_0.5')
	Dup_del('kodo_hetero_lt_0.25','No_dup_kodo_hetero_lt_0.25')
	Dup_del('kodo_min_al_freq_gt_0.05','No_dup_kodo_min_al_freq_gt_0.05')
	Dup_del('little_miss_lt_0.5','No_dup_little_miss_lt_0.5')
	Dup_del('little_hetero_lt_0.25','No_dup_little_hetero_lt_0.25')
	Dup_del('little_min_al_freq_gt_0.05','No_dup_little_min_al_freq_gt_0.05')
	Dup_del('proso_miss_lt_0.5','No_dup_proso_miss_lt_0.5')
	Dup_del('proso_hetero_lt_0.2','No_dup_proso_hetero_lt_0.2')
	Dup_del('proso_min_al_freq_gt_0.05','No_dup_proso_min_al_freq_gt_0.05')
Then use the following script to get filtered hapmaps that will take out all of the "bad site"
	#Set your working directory to where every you downloaded your files too
	workdir='/home/mjohnson/Desktop/Minor_millet_code_for_publication/'
	#This is to take the filtered files and only return sites that pass all three filters(Minor allele freq,prop. hetero, and prop miss.)
	#end the good row rows in the hap. map files with all of that info
	def HMP_Filter(hetero,min_al_freq, missing,HMP_file,new_file_name):
	    a=open(workdir + hetero+ ".txt", "r")
	    b=open(workdir+ min_al_freq + ".txt", "r")
	    c=open(workdir+ missing + ".txt", "r")
	    g=open(workdir + HMP_file , "r")
	    het=set(a)
	    min_al=set(b) 
	    miss=set(c)
	    #min_al is where the filtered site names are held
	    het.intersection_update(min_al)
	    min_al.intersection_update(het)
	    miss.intersection_update(min_al)
	    min_al.intersection_update(miss)
	    #This part of code is used to show if all fitlered sites are found in hap map file 
	    values=list()
	    min_al_list=list()
	    for sites in min_al:
		min_al_list.append(sites.strip("\n"))
	    for line in g:
		h=line.strip("\n").split("\t")
		values.append(h[0])
	    new_values=set(values)
	    new_values.intersection_update(set(min_al_list))
	    print(len(min_al),"This is the number of sites that passed through the three filter")
	    print(len(new_values),"This shows how many of the sites that passed the filters were found in the hapmap file")
	    if len(min_al)!= len(new_values):
		print("WARNING these numbers should be equal, check data integrity")
	    #this section is to filter hapmap files so we are left with just the information of the filtered sites
	    p=open(workdir + HMP_file , "r")
	    header=p.readline()
	    filtered_rows=list()
	    for rows in p:
		site=rows.strip("\n").split("\t")[0]#what does this [0] do? makes it only read first collumn i think
		if site in min_al_list:
		    filtered_rows.append(rows)
	    o=filtered_rows
	    h=open (workdir + new_file_name+ ".txt", "w")         
	    h.write(header)
	    h.writelines(o)
	    h.close()

	HMP_Filter("No_dup_kodo_hetero_lt_0.25","No_dup_kodo_min_al_freq_gt_0.05","No_dup_kodo_miss_lt_0.5","HapMaps/Raw_hmp/kodo.raw.hmp.txt","HapMaps/Filtered_kodo_hmp")
	HMP_Filter("No_dup_little_hetero_lt_0.25","No_dup_little_min_al_freq_gt_0.05","No_dup_little_miss_lt_0.5","HapMaps/Raw_hmp/little.raw.hmp.txt","HapMaps/Filtered_little_hmp")
	HMP_Filter("No_dup_proso_hetero_lt_0.2","No_dup_proso_min_al_freq_gt_0.05","No_dup_proso_miss_lt_0.5","HapMaps/Raw_hmp/proso.raw.hmp.txt","HapMaps/Filtered_proso_hmp")

Now you need to take the filtered hapmap files and get the TAXA GENO SUMMARY file to be able to filter the taxa
This is done using the following scripts:

	#You need to get a Taxa geno summary from tassel using the Filtered HMPs
	#Set your working directory to where every you downloaded your files too
	workdir='/home/mjohnson/Desktop/Minor_millet_code_for_publication/'
	#This filters out good taxa and writes out the taxa names to a file, those files might be able to be used
	#in the pandas program to delete the bad collumns from the hapmap file
	def Filter_function(folder,file_name,column_name,min_number,max_number,new_file_name):
	    f=open(workdir + folder + "/"  + file_name , "r")
	    column=-1
	    values=list()
	    for line in f:
		x=line.strip("\n").split("\t")
		if column==-1:
		    column=x.index(column_name)
		    column2=x.index("Taxa Name")
		y=(x[column])
		q=(x[column2])
		values.append([y,q])
	    filtered=list()
	    for score,name in values:
		if score!=column_name and float(score) >= min_number and float(score)<= max_number:
		    filtered.append(name+'\n')
	    o=filtered
	    h=open (workdir + new_file_name+ ".txt", "w")        
	    h.writelines(o)
	    h.close()
	Filter_function("Site_summaries","Kodo_filtered_taxa_sum.txt","Proportion Missing",0.5,1.0,"kodo_bad_taxa_miss_gt_0.5")
	Filter_function("Site_summaries","Little_filtered_taxa_sum.txt","Proportion Missing",0.5,1.0,"little_bad_taxa_miss_gt_0.5")
	Filter_function("Site_summaries","Proso_filtered_taxa_sum.txt","Proportion Missing",0.5,1.0,"proso_bad_taxa_miss_gt_0.5")   


	#Set your working directory to where every you downloaded your files too
	workdir='/home/mjohnson/Desktop/Minor_millet_code_for_publication/'
	#This uses pandas to delete columns from a hpm file that are from taxa with too much proportion missing
	def Taxa_Filter(Filtered_hmp, bad_taxa_files, new_files):
	    import pandas as pd
	    taxa_df=pd.read_table(workdir + Filtered_hmp + ".txt")
	    p=open(workdir + bad_taxa_files + ".txt", 'r')
	    bad_taxa=list()
	    for line in p:
		x=line.strip("\n")
		bad_taxa.append(x)
	    taxa_df.drop(bad_taxa, axis=1, inplace=True)

	    taxa_df.to_csv(workdir + new_files +'.txt', index=False, sep="\t")
	    print('Done')
	    
	    
	Taxa_Filter('HapMaps/Filtered_kodo_hmp','kodo_bad_taxa_miss_gt_0.5','HapMaps/Final_filt_kodo.hmp')
	Taxa_Filter('HapMaps/Filtered_little_hmp','little_bad_taxa_miss_gt_0.5','HapMaps/Final_filt_little.hmp')
	Taxa_Filter('HapMaps/Filtered_proso_hmp','proso_bad_taxa_miss_gt_0.5','HapMaps/Final_filt_proso.hmp')


Now you have the final filtered hapmaps, the previous version of tassel i had seems like it would automatically save
a hapmap as a diploid so that is what happened the first time i ran this through. When saved as a diploid hapmap file
the plink file made from those hapmap files will have a difference in the ped files then the plink files made from 
the haploid hapmap files. The heterozygous sites in the diploid file will be put into the ped in for examples: AT order
but in the haploid hapmap file when that is saved a plink the ped file will save the heterozygous site as: TA, so it 
inverts the call, this was causing issues with reproducability in the calling of the populations.

So after that was done the plink files from tassel need to have the .plk removed from them to be able to work with the 
plink program. After running the plink program run it through faststructure, i used seed=100

Then use chooseK and that should get you consistent results. 
