## Supplementary material for "Genome-wide Population Structure Analyses of Three Minor Millets: Kodo Millet, Little Millet, and Proso Millet": AltSplitstree.sh

```
#!/bin/bash

AltSplit="/home/mjohnson/Desktop/bioinfor_software/AlterSplitstreeOutput.py"
#kodo
python3 $AltSplit -i "/home/mjohnson/Desktop/Millet_Files/final_analysis/trees/kodo.svg" \
 -k "/home/mjohnson/Desktop/Millet_Files/final_analysis/trees/pop_info/kodo_7meanQ_keyfile_gt0.6.txt"\
 -o"/home/mjohnson/Desktop/Millet_Files/final_analysis/trees/color_kodo.svg" -n "unknown"\
 -c "Populations" --colorize

#little
python3 $AltSplit -i "/home/mjohnson/Desktop/Millet_Files/final_analysis/trees/little.svg" \
 -k "/home/mjohnson/Desktop/Millet_Files/final_analysis/trees/pop_info/little_8meanQ_keyfile_gt0.6.txt"\
 -o"/home/mjohnson/Desktop/Millet_Files/final_analysis/trees/color_little.svg" -n "unknown"\
 -c "Populations" --colorize

#proso
python3 $AltSplit -i "/home/mjohnson/Desktop/Millet_Files/final_analysis/trees/proso.svg" \
 -k "/home/mjohnson/Desktop/Millet_Files/final_analysis/trees/pop_info/proso_8meanQ_keyfile_gt0.6.txt"\
 -o"/home/mjohnson/Desktop/Millet_Files/final_analysis/trees/color_proso.svg" -n "unknown"\
 -c "Populations" --colorize
```
