## Supplementary material for "Genome-wide Population Structure Analyses of Three Minor Millets: Kodo Millet, Little Millet, and Proso Millet": choose_millet.sh

```
#!/bin/bash
#exec > >(tee -ia millet_choose.log)
choose="python2 /home/mjohnson/Desktop/bioinfor_software/fastStructure/chooseK.py"


for millet in kodo little proso  ; do
 

$choose --input="/home/mjohnson/Desktop/Minor_millet_Filter_pipline_code/fastStruct/"$millet"FS_genotypes_output" \
|tee ${millet}_choose.txt

done
```
