## Supplementary material for "Genome-wide Population Structure Analyses of Three Minor Millets: Kodo Millet, Little Millet, and Proso Millet": distruct_millet.sh

```
#!/bin/bash

#looks at the different files generated from fast structure to say which of the population numbers is best

distruct="python2 /home/mjohnson/Desktop/bioinfor_software/fastStructure/distruct.py"


for millet in kodo little proso  ; do
for K in `seq 1 12` ; do
$distruct -K $K --input="/home/mjohnson/Desktop/Millet_Files/final_analysis/fastStruct/"$millet"FS_genotypes_output" \
--output="/home/mjohnson/Desktop/Millet_Files/final_analysis/distruct/distruct_results_$millet.$K.svg"


done
done
```
