## Supplementary material for "Genome-wide Population Structure Analyses of Three Minor Millets: Kodo Millet, Little Millet, and Proso Millet": fastStructure_millet.sh

```
#!/bin/bash

fastStruc="python2 /home/mjohnson/Desktop/bioinfor_software/fastStructure/structure.py"

#creates the file/overwrites file if anything existed before, this will make sure you have a clean file with every rerun

echo > faststructure_millet_analysis.txt
#loop in bash, seq tells it to run 1-10, do starts loop, done finishes loop
for K in `seq 1 15` ; do 
#these analyses will take raw genotype data in a plink bed format and give a population analysis 

	echo $fastStruc -K $K --input="/home/mjohnson/Desktop/Minor_millet_Filter_pipline_code/Plink/kodo" \
--output="/home/mjohnson/Desktop/Minor_millet_Filter_pipline_code/fastStruct/kodoFS_genotypes_output" --full --seed=100 >>faststructure_millet_analysis.txt 

	echo $fastStruc -K $K --input="/home/mjohnson/Desktop/Minor_millet_Filter_pipline_code/Plink/little" \
--output="/home/mjohnson/Desktop/Minor_millet_Filter_pipline_code/fastStruct/littleFS_genotypes_output" --full --seed=100 >>faststructure_millet_analysis.txt 

	echo $fastStruc -K $K --input="/home/mjohnson/Desktop/Minor_millet_code_for_publication/Plink/proso" \
--output="/home/mjohnson/Desktop/Minor_millet_Filter_pipline_code/fastStruct/prosoFS_genotypes_output" --full --seed=100 >>faststructure_millet_analysis.txt 


done

#this reads out files to a text and then pipes it to parallel for processing

cat faststructure_millet_analysis.txt | parallel --progress
```
