## Supplementary material for "Genome-wide Population Structure Analyses of Three Minor Millets: Kodo Millet, Little Millet, and Proso Millet": Plink_millet.sh

```
#!/bin/bash

Plink="/home/mjohnson/Desktop/bioinfor_software/Plink/plink"

#this will take .ped and .map files from tassel and make .bed files that i need to use fastStructure

$Plink --noweb --file "/home/mjohnson/Desktop/Minor_millet_Filter_pipline_code/Plink/kodo" --make-bed --out /home/mjohnson/Desktop/Minor_millet_code_for_publication/Plink/kodo
$Plink --noweb --file "/home/mjohnson/Desktop/Minor_millet_Filter_pipline_code/Plink/little" --make-bed --out /home/mjohnson/Desktop/Minor_millet_code_for_publication/Plink/little
$Plink --noweb --file "/home/mjohnson/Desktop/Minor_millet_Filter_pipline_code/Plink/proso" --make-bed --out /home/mjohnson/Desktop/Minor_millet_code_for_publication/Plink/proso

#you need to delete the .plk part of the file name that is there when you export from tassel
```
