## Supplementary material for "Genome-wide Population Structure Analyses of Three Minor Millets: Kodo Millet, Little Millet, and Proso Millet": Maps.ipynb

```
# %% [code]
from mpl_toolkits.basemap import Basemap
import numpy as np
import matplotlib.pyplot as plt
import pandas as pd
#llcrnrlat,llcrnrlon,urcrnrlat,urcrnrlon
# are the lat/lon values of the lower left and upper right corners
# of the map.
# lat_ts is the latitude of true scale.
# resolution = 'c' means use crude resolution coastlines.
gps_df=pd.read_csv('/home/mjohnson/Desktop/Millet_Files/final_analysis/MDS/Location/Little_GPS_and_pops.csv',delimiter="\t")
m = Basemap(projection='merc',llcrnrlat=0,urcrnrlat=30,\
            llcrnrlon=50,urcrnrlon=100,lat_ts=20,resolution='c')
m.drawcoastlines()
m.fillcontinents(color='lightgrey',lake_color='white')
m.drawmapboundary(fill_color='white')
m.drawcountries()
lat = gps_df['Lat'].values
lon = gps_df['Long'].values
lons, lats = m(lon, lat)
# plot points as red dots
m.scatter(lons, lats, marker = 'o', color=gps_df['color'], zorder=5)
plt.title("Little Millet")
plt.savefig('/home/mjohnson/Desktop/Millet_Files/final_analysis/MDS/Location/Little_gps.svg')
plt.show()


# %% [code]
from mpl_toolkits.basemap import Basemap
import numpy as np
import matplotlib.pyplot as plt
import pandas as pd
#llcrnrlat,llcrnrlon,urcrnrlat,urcrnrlon
# are the lat/lon values of the lower left and upper right corners
# of the map.
# lat_ts is the latitude of true scale.
# resolution = 'c' means use crude resolution coastlines.
gps_df=pd.read_csv('/home/mjohnson/Desktop/Millet_Files/final_analysis/MDS/Location/Kodo_GPS_and_pops.csv',delimiter="\t")
m = Basemap(projection='merc',llcrnrlat=0,urcrnrlat=30,\
            llcrnrlon=50,urcrnrlon=100,lat_ts=20,resolution='c')
m.drawcoastlines()
m.fillcontinents(color='lightgrey',lake_color='white')
m.drawmapboundary(fill_color='white')
m.drawcountries()
lat = gps_df['Lat'].values
lon = gps_df['Long'].values
lons, lats = m(lon, lat)
# plot points as red dots
m.scatter(lons, lats, marker = 'o', color=gps_df['color'], zorder=5)
plt.title("Kodo Millet")
plt.savefig('/home/mjohnson/Desktop/Millet_Files/final_analysis/MDS/Location/Kodo_gps.svg')
plt.show()

# %% [code]
```
