## Supplementary material for "Genome-wide Population Structure Analyses of Three Minor Millets: Kodo Millet, Little Millet, and Proso Millet": 9_RunTasselGbs.sh

```
#! /bin/bash

TASSEL5="perl $HOME/jgwall/Software/TASSEL/tassel-5-standalone/run_pipeline.pl -Xms10g -Xmx50g"

flowcells=$HOME/Projects/0_RawData/GBS/ICRISAT_SmallMillets
workdir=9_TasselGBS
min_tag_count=10
enzyme="PstI"

if [ ! -e $workdir ]; then mkdir $workdir; fi

# # Make master keyfiles
allkeys=$flowcells/*.keys.txt
cat $allkeys | head -n 1 > $workdir/proso.keys.txt
cat $allkeys | head -n 1 > $workdir/kodo.keys.txt
cat $allkeys | head -n 1 > $workdir/little.keys.txt
grep -P --no-filename "Panicum\tmiliaceum" $allkeys >> $workdir/proso.keys.txt
grep -P --no-filename "Panicum\tsumatrense" $allkeys >> $workdir/little.keys.txt
grep -P --no-filename "Paspalum\tscrobiculatum" $allkeys >> $workdir/kodo.keys.txt

# go through species and do GBS analysis
for species in kodo proso little; do
  keyfile=$workdir/$species.keys.txt
  database=$workdir/$species.gbs_database.sql
  udir=$workdir/${species}_uneak
  if [ ! -e $udir ]; then mkdir $udir; fi

  # Make initial database
  $TASSEL5 -GBSSeqToTagDBPlugin -i $flowcells -k $keyfile -db $database -e $enzyme -c $min_tag_count

#   Export tag counts, run UNEAK on them, and convert to a SAM file of pseudo-positions
  $TASSEL5 -GetTagSequenceFromDBPlugin -db $database -o $udir/9a_$species.tags.txt
  python3 9b_ConvertTagSeqsToCountFile.py -i $udir/9a_$species.tags.txt -o $udir/9a_$species.tags.cnt.txt --pad 64 --count $min_tag_count
  $TASSEL5 -TextToBinaryPlugin -i $udir/9a_$species.tags.cnt.txt -o $udir/9a_$species.tags.cnt -t TagCounts
  $TASSEL5 -UTagCountToTagPairPlugin -inputFile $udir/9a_$species.tags.cnt -outputFile $udir/9a_$species.tags.tps
  $TASSEL5 -UTagPairToTOPMPlugin -input $udir/9a_$species.tags.tps -toText $udir/9a_$species.topm.txt
  python3 9c_ConvertTopmToFakeSam.py -i $udir/9a_$species.topm.txt -o $udir/9c_$species.topm.sam   
  
  # Export tags and map to genome (=50k unassembled contigs in this case)
  $TASSEL5 -SAMToGBSdbPlugin -i $udir/9c_$species.topm.sam -db $database

  # Run Discovery SNP Caller
  $TASSEL5 -DiscoverySNPCallerPluginV2 -db $database
  
  # Call SNPs
  $TASSEL5 -ProductionSNPCallerPluginV2 -db $database -e $enzyme -i $flowcells -k $keyfile -o $workdir/$species.raw.h5  # For some reason the export-to-VCF is not working
  $TASSEL5 -h5 $workdir/$species.raw.h5 -export $workdir/$species.raw.hmp.txt.gz -exportType Hapmap
#   break
done
```
