## Supplementary material for "Genome-wide Population Structure Analyses of Three Minor Millets: Kodo Millet, Little Millet, and Proso Millet": 9b_ConvertTagSeqsToCountFile.py

```
__author__ = 'jgwall'

import argparse

debug = False


def main():
    args = parse_args()

    # Get count of tags
    print("Counting tags in",args.infile)
    ntags=0
    for line in open(args.infile):
        ntags+=1
    ntags-=1    # To remove header
    print("\tFound",ntags,"tags")

    # Write out new file
    IN = open(args.infile, "r")
    IN.readline()   # Remove header
    OUT = open(args.outfile, "w")
    OUT.write(str(ntags) + "\t" + str(round(args.pad/32)) + "\n")

    n=0
    for line in IN:
        n+=1
        if debug and n > 1000: break
        tag = line.strip()
        taglen = len(tag)
        # Add padding
        if len(tag) < args.pad:
            tag += args.pad_nucleotide * (args.pad - len(tag))
        OUT.write(tag + "\t" + str(taglen) + "\t" + str(args.count) + "\n")
    IN.close()
    OUT.close()
    print("Wrote",n,"tags to",args.outfile)


def parse_args():
    parser = argparse.ArgumentParser()
    parser.add_argument("-i", "--infile")
    parser.add_argument("-o", "--outfile")
    parser.add_argument("-p", "--pad", type=int, default=64, help="Pad tags to this value")
    parser.add_argument("-n", "--pad-nucleotide", default="A", help="Nucleotide to pad with")
    parser.add_argument("-c", "--count", type=int, default=1, help="Fake tag count to output")
    parser.add_argument("--debug", default=False, action="store_true")
    args = parser.parse_args()

    # Handle debug flag
    global debug
    debug = args.debug

    return parser.parse_args()


if __name__ == '__main__': main()
```
