## Supplementary material for "Genome-wide Population Structure Analyses of Three Minor Millets: Kodo Millet, Little Millet, and Proso Millet": 9c_ConvertTopmToFakeSam.py

```
__author__ = 'jgwall'

import argparse

debug = False

# SAM stuff
sam_default = ["unknown", 0, "unknown", 0, 0, "*", "*", "8", 0, "*", 0]
nameID, flagID, refID, posID, lenID, seqID = 0,1,2,3,8,9

def main():
    args = parse_args()

    # Parse TOPM header
    TOPM = open(args.infile, "r")
    header=TOPM.readline().strip().split('\t')
    print("Parsing",args.infile,"which has",header[0],"tags")

    # Parse subsequent lines
    samdata=list()
    chroms=dict()   # dictionary of chromosome->position so as to find chromosome sizes
    n=0
    for line in TOPM:
        n+=1
        data=line.strip().split('\t')
        tag, taglength, mychrom, mystart, mystop = data[0], int(data[1]), data[4], int(data[5]), int(data[6])
        if len(tag) > taglength: tag=tag[:taglength]    # Trim tag to non-padded bases

        # Determine chromosome length by storing the largest stop position encountered so far, including if is a new chromosome
        if (mychrom not in chroms) or (mystop > chroms[mychrom]):
            chroms[mychrom] = mystop

        # Make SAM data line
        mysam = sam_default.copy()
        mysam[nameID] = "tagSeq=" + tag
        mysam[refID] = mychrom
        mysam[posID] = mystart
        mysam[lenID] = taglength
        mysam[seqID] = tag
        mysam = "\t".join([str(x) for x in mysam]) + "\n"
        samdata.append(mysam)
    TOPM.close()
    print("\tParsed",n,"tags. Outputting to",args.outfile)


    # Make SAM header
    SAM = open(args.outfile, "w")
    SAM.write("@HD\n")
    SAM.write("@CO Fake SAM file produced to feel into the TASSEL GBSv2 SNP-calling pipeline\n")

    # Add chromsome lengths
    for mychrom in sorted(chroms.keys()):
        SAM.write("@SQ SN:" + mychrom + " LN:" + str(chroms[mychrom] + 10) + "\n")

    # Add tag data
    SAM.writelines(samdata)
    SAM.close()

def parse_args():
    parser = argparse.ArgumentParser()
    parser.add_argument("-i", "--infile", help="TASSEL TOPM file in text format")
    parser.add_argument("-o", "--outfile", help="TASSEL-readable SAM file (probably wouldn't work for SAMtools)")
    parser.add_argument("--debug", default=False, action="store_true")
    args = parser.parse_args()

    # Handle debug flag
    global debug
    debug = args.debug

    return parser.parse_args()


if __name__ == '__main__': main()
```
