## Supplementary material for "Genome-wide Population Structure Analyses of Three Minor Millets: Kodo Millet, Little Millet, and Proso Millet": AlterSplitstreeOutput.py

```
__author__ = 'jgw87'
"""
Relabel an SVG graphic made by SplitsTree4 according to data given in an external file
    -i SVG file made by SplitsTree4
    -o Output file name
    -k Keyfile with pop information; should have a column with "sample" corresponding to existing labels
    -c Column name to pull new label from
    -n Null color code - which code value in the keyfile should not be colorized
    --colorize  Whether to color the labels based on their new values
    --keep_labels   Whenther to keep the original labels (if only want to colorize)
"""

import argparse
import math
import matplotlib.cm as cm
import math
import numpy as np
import pandas as pd
import re


def main():
    args = parse_args()

    print("Altering SVG labels in",args.infile)
    key = get_change_key(args.keyfile, args.colname)
    colorkey = None
    if args.colorize:
        colorkey = get_color_key(key)
    alter_svg(args.infile, key, args.outfile, colorkey, args.keep_labels, args.null_color)

def parse_args():
    parser = argparse.ArgumentParser()
    parser.add_argument("-i", "--infile")
    parser.add_argument("-c", "--colname")
    parser.add_argument("-o", "--outfile")
    parser.add_argument("-k", "--keyfile")
    parser.add_argument("-n", "--null_color")
    parser.add_argument("--colorize", default=False, action='store_true')
    parser.add_argument("--keep_labels", default=False, action='store_true')
    args = parser.parse_args()
    return args

def get_change_key(keyfile, colname):
    print("\tMaking substitution key")
    data = pd.read_csv(keyfile, sep="\t")

    key=dict()
    for sample, newlabel in zip(data['sample'], data[colname]):
        sample=str(sample)        
        key[sample]=str(newlabel)
        if key[sample] == "nan":
            key[sample] = "Unknown"
    return key

def alter_svg(infile, key, outfile, colorkey, keeplabels, nullcolor):
    print("\tAltering sample names to origin codes")
    OUT = open(outfile, "w")
    textfinder=re.compile(r"<text")      #this might not be reading my svg file properly 
    namefinder=re.compile(r">(.+)</text")

    #Go through and scan lines for samples
    found=0
    for line in open(infile, "r"):
        matcher = namefinder.search(line)
        if matcher is None: #If line doens't have the pattern for a sample name, write and continue
            OUT.write(line)
            continue

        found+=1
        sample=matcher.group(1)
        if not keeplabels:
            if sample in key:
                label = key[sample]
            else: label="Unknown"
            newline=namefinder.sub(repl=">" + label + "</text>", string=line)
        else: newline = line

        if (colorkey is not None) and (sample in key) and (key[sample] != nullcolor):
            matcher = namefinder.search(newline)
            color = colorkey[key[sample]]
            color = [str(math.floor(x*255)) for x in color[:3]]
            colortext=" fill=\"rgb(" + ",".join(color) + ")\"" + matcher.group()
            newline=namefinder.sub(repl=colortext, string=newline)
        OUT.write(newline)
    OUT.close()
    print("\t\tFound",found,"samples to alter labels of")

def get_color_key(key):
    print("\tMaking color key")
    values = sorted(np.unique(list(key.values())))
    colorspace = cm.rainbow(np.linspace(0, 1, len(values)))

    #Match color to species
    colorkey=dict()
    for i in range(len(values)):
        colorkey[values[i]]=colorspace[i]
    return colorkey


if __name__ == "__main__":
    main()
```
