## Supplementary material for "Genome-wide Population Structure Analyses of Three Minor Millets: Kodo Millet, Little Millet, and Proso Millet": Millet_make_csv_for_pop_structure.ipynb

```
# %% [code]
#takes meanQ files from FastStuc output and makes them into csv files instead of one row

def make_csv(input_file, output_file):
    p=open("/home/mjohnson/Desktop/Millet_Files/final_analysis/trees/pop_info/" + input_file)
    csv=list()
    for row in p:
        row=row.replace('  ','\t') #you need to capture the value or else it will dissapear, that is why you need row=
        csv.append(row)
    o=csv
    h=open ("/home/mjohnson/Desktop/" +output_file , "w")        
    h.writelines(o)
    h.close()    
#make_csv('kodo_7.meanQ','kodo_mod_7.meanQ')
#make_csv('little_8.meanQ','little_mod_8.meanQ')
make_csv('proso_9.meanQ','proso_mod_9.meanQ')

# %% [code]
#i can maybe modify this filter script to filter the meanQ files to make a file with an assigned population
def make_keyfile(i_file,o_file):
    f=open("/home/mjohnson/Desktop/Millet_Files/final_analysis/trees/pop_info/"+ i_file , "r")
    pop_values=list()
    #this next step give a list of pop_1-pop_7
    col_names = f.readline().strip().split(',')[1:]
    paired=list()

    taxa_pops=list()

    for line in f:
        x=line.strip('\n').split(',')
        taxon=x[0]
        x=x[1:] #this has the list ignore the first values, so taxa names ignored 
        pops="unknown"
        for i in range(len(x)):
        #x[i] is taking the value number 'i' in the list of x
            if float(x[i]) > 0.85:
        #this will put the name of the population that is geater than .5, and if none are it will put 'unknown'
                pops=col_names[i]
            taxon=taxon[:10]
        taxon2=taxon.replace(':','_')
        taxa_pops.append(taxon2+'\t'+ pops+ '\n')    
    taxa_pops.insert(0,'sample\tPopulations\n')
    o=taxa_pops
    h=open ("/home/mjohnson/Desktop/Millet_Files/final_analysis/trees/pop_info/"+o_file, "w")        
    h.writelines(o)
    h.close()

#make_keyfile('kodo_mod_7.meanQ','kodo_7meanQ_keyfile_gt0.6.txt')
#make_keyfile('little_mod_8.meanQ','little_8meanQ_keyfile_gt0.6.txt')
make_keyfile('proso_mod_6.meanQ','proso_6meanQ_keyfile_gt0.85.txt')


# %% [markdown]
```
