## Supplementary material for "Genome-wide Population Structure Analyses of Three Minor Millets: Kodo Millet, Little Millet, and Proso Millet": Principle_components_scatter_plot.ipynb

```
# %% [code]
import numpy as np
import matplotlib.pyplot as plt
import pandas as pd
#i was plotting with pandas and not matplotlib at first and that was casuing problems
#the colors are done using the hexcode, which was attained from the fill section on inkscape
pc_df=pd.read_table("/home/mjohnson/Desktop/Millet_Files/MDS/MDS_PC_kodo.txt")
color_key=dict(Pop_1='#7f00ff', Pop_2='#497af8', Pop_3='#40d2ea', Pop_4='#f12133', Pop_5='#197b47', Pop_6='#ecc76e', Pop_7='#f047b0', unknown='grey')
pop_color=list()
for pop in pc_df['Populations']:
    pop_color.append(color_key[pop])
plt.scatter(x=pc_df['PC1'], y=pc_df['PC2'], color=pop_color, s=40)
plt.xlabel('PC1')
plt.ylabel('PC2')
plt.title('Kodo Millet PC Analysis')
plt.savefig("/home/mjohnson/Desktop/Millet_Files/MDS/Kodo_PC_analysis.svg")
plt.show()


# %% [code]
import numpy as np
import matplotlib.pyplot as plt
import pandas as pd
#i was plotting with pandas and not matplotlib at first and that was casuing problems
#the colors are done using the hexcode, which was attained from the fill section on inkscape
pc_df=pd.read_table("/home/mjohnson/Desktop/Millet_Files/MDS/MDS_PC_proso.txt")
color_key=dict(Pop_1='#7f00ff', Pop_2='#497af8', Pop_3='#40d2ea', Pop_4='#f12133', Pop_5='#197b47', Pop_6='#ecc76e', Pop_7='#f047b0', Pop_8='#fe7f29', unknown='grey')
pop_color=list()
for pop in pc_df['Populations']:
    pop_color.append(color_key[pop])
plt.scatter(x=pc_df['PC1'], y=pc_df['PC2'], color=pop_color, s=40)
plt.xlabel('PC1')
plt.ylabel('PC2')
plt.title('Proso Millet PC Analysis')
plt.savefig("/home/mjohnson/Desktop/Millet_Files/MDS/Proso_PC_analysis.svg")
plt.show()

# %% [code]
import numpy as np
import matplotlib.pyplot as plt
import pandas as pd
#i was plotting with pandas and not matplotlib at first and that was casuing problems
#the colors are done using the hexcode, which was attained from the fill section on inkscape
pc_df=pd.read_table("/home/mjohnson/Desktop/Millet_Files/MDS/MDS_PC_little.txt")
color_key=dict(Pop_1='#7f00ff', Pop_2='#497af8', Pop_3='#40d2ea', Pop_4='#f12133', Pop_5='#197b47', Pop_6='#ecc76e', Pop_7='#f047b0', Pop_8='#fe7f29', unknown='grey')
pop_color=list()
for pop in pc_df['Populations']:
    pop_color.append(color_key[pop])
plt.scatter(x=pc_df['PC1'], y=pc_df['PC2'], color=pop_color, s=40)
plt.xlabel('PC1')
plt.ylabel('PC2')
plt.title('Little Millet PC Analysis')
plt.savefig("/home/mjohnson/Desktop/Millet_Files/MDS/Little_PC_analysis.svg")
plt.show()
```
