## Supplementary material for "Genome-wide Population Structure Analyses of Three Minor Millets: Kodo Millet, Little Millet, and Proso Millet": Minor_Millet_SupplementalFiles_Readme

This goes over some of the infomration is found in the supplemental files.

Minor_millet_Filter_pipeline_code
This gives all the files and code needed to replicate the filtereing of the raw hapmaps and the calling of the populations
----See read me in this folder for more in-depth information

Maps
This shows the millets plotted by known geographic locations and colored by population 

Phenotype_data
This take the information found in Upadhyaya et al., 2014 supplemental data (see paper for complete reference) and filters it down to just the accessions that we had genotype data for and looks only at the  phenotype data. 

PC_analysis
This folder containes two principal coordinate plots showing that race is correlated with population structure.

Key_Files
These files contain the information on the samples that had GBS preformed on them

HapMaps
This folder contains the raw SNP data on the three millet species and the filtered SNP data. To see filtering
parameters see the materials and methods section of the paper or look through the code folder in this 
supplemental data section. 

Code: The .ipynb file format that can be opened with jupyter notebook 
-Principle_components_scatter_plot: This shows the code used to generate the scatterplot figures
-Millet_make_csv_for_pop_structure: This takes the meanQ files from FastStructure and alters it into a csv
and takes the assigned population to create a keyfile
-AtlerSplitstreeOutput.py: Code developed by jason wallace to alter an svg file from splitstree4 to show 
population information instead of accession names
