## Supplementary figures and images for "Genome-wide Population Structure Analyses of Three Minor Millets: Kodo Millet, Little Millet, and Proso Millet"

### Little_gps.png

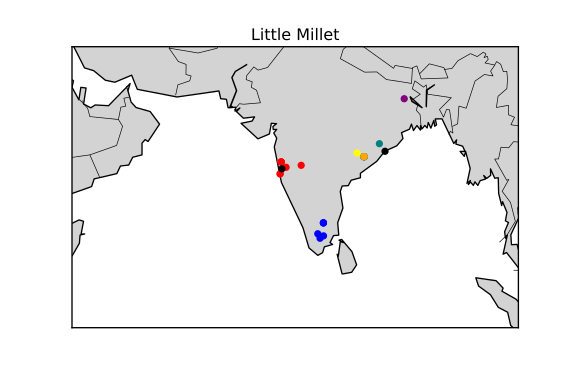
